# Detection and electron microscopic study of thick cross-striated linear fibrils in mammalian cell nuclei

**DOI:** 10.1101/2024.10.18.619048

**Authors:** Mark I. Mosevitsky

**Affiliations:** Institute of Macromolecular Compounds. Saint-Petersburg, Russian Federation

**Keywords:** cell nucleus architecture, nuclear matrix, nuclear matrix fibrils, chromatin and nuclear matrix contacts, preservation of nuclear matrix elements in mitosis

## Abstract

Using an electron microscope, cross-striated fibrils with an axial repeat of about 65 nm were detected in the cell nuclei of rat and calf tissues. The nuclei were purified and processed using a “complete medium” (CM) simulating the composition of salts of the intranuclear medium. These fibrils are not the filaments of the nuclear matrix described in the literature. In particular, they are thicker and not branched. The axial repeat in these fibrils is similar to that in extracellular collagen fibrils. Therefore, in this work, the main efforts were aimed at demonstrating the nuclear origin of the observed fibrils. Their presence in the material of nuclei destroyed by ultrasound, their contact with isolated nucleoli and their presence in residual nuclei are shown on samples prepared for electron microscopy in the form of “whole mounts”. The sites of attachment of chromatin and of nuclear matrix network to thick cross-striated fibrils were detected. As axial components of condensed chromosomes, thick cross-striated fibrils are preserved during mitosis. Probably, their contacts with chromatin and with elements of nuclear matrix network are also preserved providing reproduction of the internal structure of the nuclei in daughter cells. It is logical to assume that the cross-striated fibrils perform important functions attributed to the nuclear matrix.

**Highlights:** Thick cross-striated fibrils with an axial repeat of about 65 nm are present in mammalian cell nuclei.

Areas of dense contact between thick cross-striated fibrils and chromatin are observed.

Nucleoli are attached to thick cross-striated fibrils.

Nuclear matrix network is tightly adherent to thick cross-striated fibrils.

Thick cross-striated fibrils can be retained in mitosis as axial elements of chromosomes.

## Introduction (review of the literature on the structure and functions of the nuclear matrix)

Insoluble at high ionic strength (1-2 M NaCl) components of the cell nucleus were first studied more than 60 years ago (Zbarsky, Georgiev, 1959).^1^ Subsequent studies have shown that this material is preserved after chromatin removal with the combined action of, 2 M NaCl and DNase (in some cases, endogenous digestion was used instead of DNase).

Electron microscopic studies of embedded in a polymer or resin ultrathin sections of these “residual nuclei” have demonstrated an irregular granulofibrillary network filling the entire volume of the nucleus. (Georgiev Chentsov, 1962; Berezney Coffey, 1974; Valter et al., 1984).^2,3,4^ This substance was called nuclear matrix or nuclear skeleton (scaffold). The periphery of the nuclear matrix underlying the nucltar envelop, loocs more dens then its internal part It was called lamina. So, the nuclear matrix is formed by lamina and internal matrix.

Researcher’s interest in the nuclear matrix increased dramatically after it was discovered that pulse-labeled (newly synthesized) DNA was detected in this “residual” substance (Berezney and Coffey, 1975; McCready et al., 1980),^5,6^ where RNA formation was also shown to occur (Razin and Yarovaya, 1985; Wei et al., 1999).^7,8^, An important contribution to these results was made by the use of electron microscopic autoradiography. In the rat regenerating liver, Berezney and collaborators labeled DNA with [^3^H] thymidine for a long time (60 minutes) to localize DNA fixation sites and by impulses (for 1 minute) to localize DNA replication sites (Smith et al., 1984; Berezney et al., 1995).^9,10^ After a long period of labeling, endogenous DNA cleavage was performed. Loose (not fixed) labeled DNA fragments were removed by the following extractions, including DNase activity. Thin sections of whole tissue, isolated nuclei and nuclear matrices were prepared. They were mounted on electron microscopic meshes, covered with photoemulsion and exposed for 6-8 months. Observation of the developed autoradiographs in an electron microscope showed that most of the DNA fixation sites (long-term labeling) and DNA replication sites (pulse labeling) are mainly localized in the internal nuclear matrix. Similar experiments with the same results were obtained by Pardoll et al.^11^

In other kind of experiment, synchronized Murine 3T3 cells were used. It was was syown that when during cell cycle they reached S-phase, necessary for initiation of DBA replication helicase id found assotiated with the nuclear matrix (Hesketh et al.,2015).^12^

To find the location of the RNA synthesis sites, cells HeLa were labeled with [^3^H]uridine. Then the nuclei were treated with DNase and RNase. Thin slices of residual nuclei (nuclear matrix) were prepared. Autoradiographs were prepared, as described above. Their observation in the electron microscope showed that transcription also occurs in the internal nuclear matrix (Herman et al., 1978).^13^ This result was confirmed by immunochemical localization of proteins carrying out transcription in the internal nuclear matrix (Jackson et al., 1993; Grande et al., 1997; Szentirmay and Sawadogo, 2000).^14.15.16^ Observation of fluorescently labeled nascent RNA reveals transcription in domains scattered throughout the nucleus (Wansink et al., 1993).^17^

Transformation of primary transcripts into messenger RNA (splicing) is carried out by multifunctional enzyme complexes also located in the internal nuclear matrix (Zeitlin et al., 1987; Smith et al., 1989; Jackson et al., 1993; Blencowe et al., 1994; Chabot et al., 1995;. Wagner et al., 2003; Vester et al., 2022).^14,18,19,20,21,22,23^ It has been shown that damaged DNA binds to the nuclear matrix during its repair. This means that the enzymes that carry out DNA repair are also located in the nuclear matrix (McCready and Cook, 1984; Koehler and Hanawalt, 1996).^24,25^ So, we come to the conclusion that all the main genetic processes are carried out by enzymatic complexes located in the nuclear matrix.

Evidently, nuclear matrix, being the place, where enzymes performing genetic reactions are fixed, should also ensure the contact of these enzymes with their substrate – DNA. Indeed, DNA contains the non-coding regions of 1000-3000 nucleotides long, called the scaffold/matrix attachment regions (S/MAR) with special composition rich in A-T pairs tandem repeats (Mirkovitch et al., 1984; Romig et al., 1994; Podgornaya, 2022).^26,27,28^ It was found that in the nuclear matrix these DNA regions bind to an abundant nuclear matrix protein called scaffold attachment factor A (SAF-A). This protein contains a domain (SAF-Box) that can fix S/MAR (Romig et al., 1992; Fackelmayer et al., 1994; Kipp et al., 2000).^29,30,31^

This discrete fixation of DNA in the nuclear matrix organizes chromosomal DNA (chromatin) in the form of a sequence of loops that are separate (аutonomous) replicons (Razin et al., 1986; Anachkova et al., 2005; Kitamura et al., 2006).^32,33,34^

These loops were visualized by Lemmley and his colleagues using electron microscopy (Mirkovitch et al., 1984, 1988).^26,35^

Enzyme ensembles carrying out genetic processes that are attached to the nuclear matrix should be located near the DNA fixation points (loop bases). Each loop should contain a complete set of these enzyme ensembles (Nayler et al., 1998).^36^

To perform a specific genetic function, a looped DNA passes through an appropriate ansemble of enzymes. A probable mechanism of DNA replication using a fixed enzyme ensamble (replisome) has been proposed (Mosevitsky, 1976; Alberts, 1984).^37,38^

It is important to note that the level of gene activity is determined not only by interaction with transcription factors, but also by the position of the gene in the loop, since proximity to the fixed in the nuclear matrix enhances its activity (Robinson et al., 1982).^39^ Possibly, the vicinity to the point of fixation affects the conformation of the genes. The DNA regions fixed in the nuclear matrix change according to a certain program during development and are different in different kinds of cells (Sureka et al., 2022).^40^ As a result, the composition and size of the loops and the position of the genes in them change. Consequently, the degree of activation of certain genes changes due to this epigenetic process, which also depends on the nuclear matrix (Robinson et al., 1982; Chunduri et al., 2022).^39,41^ In chromosomal DNA, in addition to the regions for strong fixation in the nuclear matrix, there are not so long sections in the intergenic regions that are used for temporary (transient) binding of the activated gene to the site of the matrix where transcribing enzymes are located, or of the damaged DNA region to the site of the matrix where the restoring ensemble is located (Chinn and Comai, 1996; Koehler and Hanawalt, 1996; Heng et al., 2004; Iarovaia et al., 2005).^25,42,,43,44^ Interestingly, RNA somehow participates in formation of temporary DNA-matrix contacts (Patriotis et al., 1990).^45^

Fixation of specific DNA regions in the nuclear matrix is one of the functions of SAF-A. The DNA (chromatin) loops formed as a result of this interaction are not conservative. DNA fixation points and, accordingly, loops vary according to certain programs in different tissues, which is one of the main epigenetic processes.

Near DNA regions recognized by SAF-A and attached to the nuclrar matrix contaih replication origin recognized by enzyme ensemble performing DNA replication. Therefore the replication fork was found also to be attached to the nuclear matrix (Aelen et al., 1983; Opstelten et al., 1989).^46,47^

The second function of SAF-A is to bind associated with chromatin long non-coding RNA (lncRNA). This interaction affects chromatin conformation and therefore can influence the gene transcription. In these interactions, SAF-A can act as oligomer (Nozawa et al., 2017; Marenda et al., 2022).^48,49^

Above only SAF-A was noted as an agent connecting chromosomal DNA and associated with chromatin non-coding RNA (RNP) with the nuclear matrix. In fact, similar functions fulfills other nuclear matrix protein SAF-B (Renz and Fackelmayer, 1996; Kipp et al., 2000; Cherney et al., 2023).^31,50,51^

It was shown that the fibrils formed by the abundant nuclear matrix proteins lamins also bind DNA (Ludérus et al., 1992, 1994; Pennarun et al., 2023)^52,53,54^ and chromatin histones (Taniura et al., 1995).^55^

Nuclear matrix proteins can be active participants in genetic processes. So, the NMP-1 protein belonging to the nuclear matrix participates in transcription as a transcription factor (Bidwell et al., 1993; Guo et al., 1995).^56,57^ The data presented above show that the nuclear matrix participates in genetic processes both as a framework on which enzyme complexes and specific regions of chromosomal DNA are fixed, and as a direct regulator of chromatin conformation and gene activity.

All these well-documented results show that detailed information on the composition and architecture of the nuclear matrix is necessary for a comprehensive study of chromatin conformations and genetic processes.

Using SDS Polyacrylamide gel electrophoresis (PAGE) and two-dimensional PAGE, a significant number of proteins was detected in the nuclear matrix (Peters and <u>Commings</u>, 1980; Peters et al., 1982; <u>Kaufmann</u>, 1989; Cupo et al., 1990; Ma et al., 1999; Bihani et al., 2023).^58,59,60,61^ Four of them with molecular weights in the range of 55-70 kDa are strongly predominant (Berezney and Coffey, 1977;. Gerace and Blobel, 1980).^62,63^ They were named lamins A, B1, B2 and C (Gerace and Blobel, 1980; Buxboim at al., 2023).^63,64^ Mutant lamins can cause different diseases called laminopathies (Broers et al., 2006; Forleo et al., 2015; Lee et al., 2016; Mosevitsky, 2022; Rootwelt-Norberg et al., 2023).^65,66,67,68,69^

Lamins A and C are coded by the same gene. Their mRNAs are formed by alternative splicing. Two lamins B1 and B2 are coded by different genes. All lamins are fibrillar proteins with similar structure (Fisher et al., 1986; McKeon et al., 1986; Stuurman et al., 1998; Herrmann., 2003; Goldberg et al., 2008; Gruenbaum and Foisner, 2015).^70,71,72,73.74,75^

*In vitro*, lamins can aggregate forming “paracrystals” that have very distinct cross-striated structure with the axial repeat of 20-27 nm (Aebi et al., 1986; Stuurman et al., 1998; Moir et al., 2000; Karabinos et al., 2003; Turgay et al., 2017; Sapra and Medalia 2021).^72,76,77,78,79,80^

On the human erythroleukemia cells with the use of immunofluorescence microscopy, it was shown that lamin A is present in the internal nuclear matrix (Neri et al., 1999).^81^

Still one critically important constituent of the internal nuclear matrix proved to be RNA. The treatment of residual nuclei (nuclear matrix) with RNase made internal nuclear matrix soluble, and it could be removed by washes. This result claims that RNA is responsible for stability of the internal nuclear matrix. However peripheral more dense area named lamina remained stable. Therefore thin sections of these structures looked in the electron microscope as rings with empty interior (Herman et al., 1978; Smith et al., 1984).^9,13^

The main components of lamina are lamins of the both groups A and B (Gerace and Blobel, 1980). These proteins form two types of fibrils consisting of lamin pairs: A and C or B1 and B2 (Goldberg et al., 2008).^74^ Connective proteins bind these fibrils to the nuclear envelope making it more rigid. Inner side of this layer of lamin fibrils contacts with the internal matrix network and with periphery of interphase chromosomes influencing their conformation (Hozak et al., 1995; Vlcek et al., 2001; Goldberg et al., 2008; Butin-Israeli et al., 2012; Xie et al., 2016; de Leeuw et al., 2018; Zheng et al., 2018).^74,82,83,84,85,86,87^

Further progress in the study of the architecture, composition and functions of the nuclear matrix was achieved when thick resinless sections and whole mounts of the residual nuclei began to apply. In these “3D” preparations, the network of long branched fibrils (filaments) with the globular inclusions as constituents of the internal nuclear matrix, as well as a peripheral lamina formed by more densely spaced fibrils, were observed in an electron microscope (Capco et al., 1982; Penman et al., 1982; Fey et al., 1986; He et al., 1990; Hozak et al., 1993, 1995; Nickerson, 2022).^82,88,89,90,91,92,93^

Different publications provide significantly different data on the properties of nuclear matrix filaments (fibrils) observed in an electron microscope.

In study of Capco et al. (1982), ^88^ diameter of the branched filaments of internal nuclear matrix ranges from 3 to 22 nm. According to He et al. (1990), ^91^ diameter of the branched “core” filaments is of about 10 nm (Fig. 1 Introd). Jackson and Cook (1988), ^94^ using the same cells (HeLa), observed in the nuclear matrix branched filaments with diameter of about 10 nm, which are not smooth, but cross-striated, with an axial repeat of 23 nm (Fig.2 Introd).

**Fig. 1 Introd.**
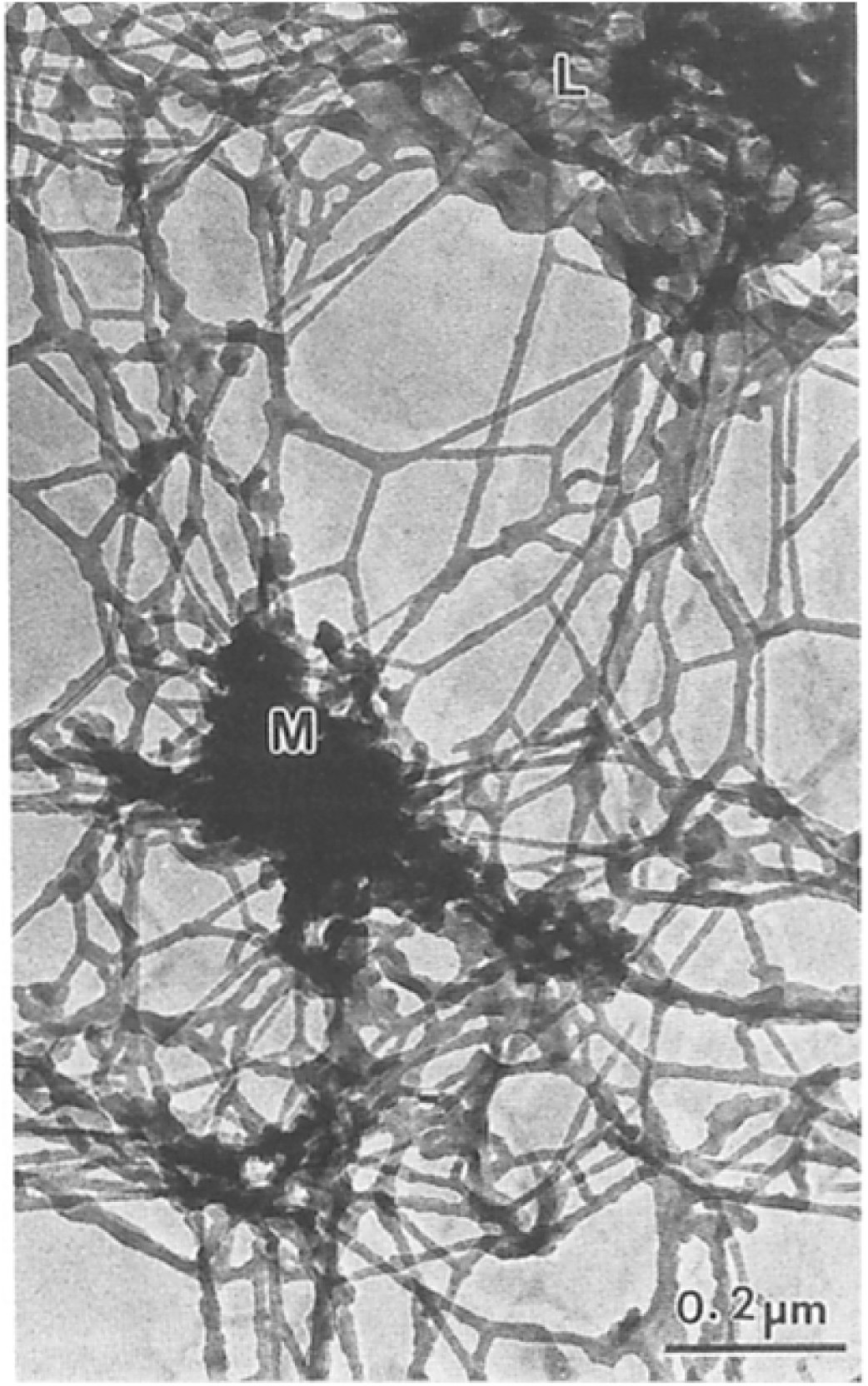
Filaments of the nuclear matrix prepared from HeLa cells. Electron microscopy (Fig.5b in He et al., 1990 ^91^, License CC-BY 4.0)

**Fig. 2 Introd.**
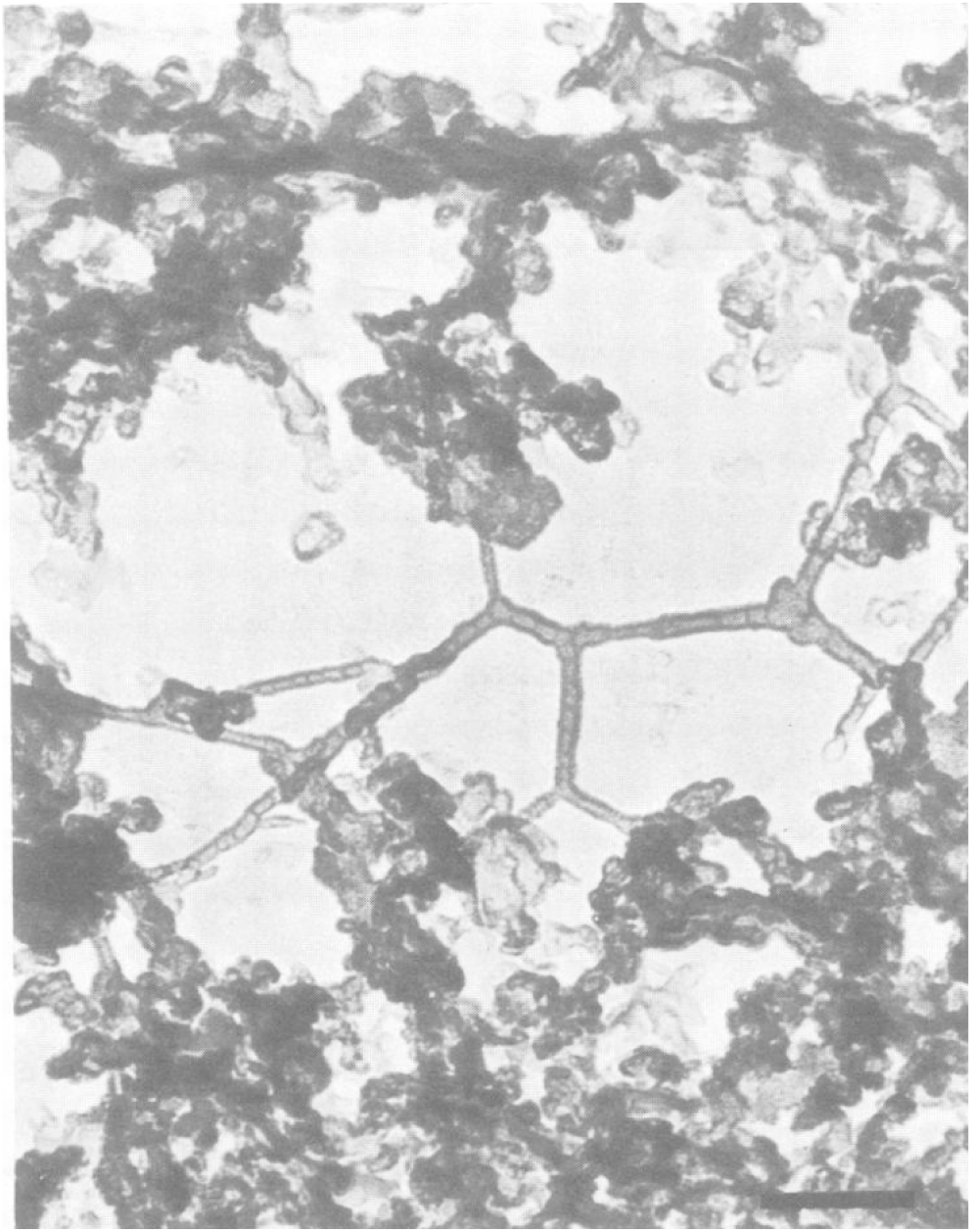
Cross-striated filaments of the nuclear matrix prepared from HeLa cells. Electron microscopy, Bar 0.1 µm. (Fig. 3c in Jackson and Cook, 1988 ^94^, License CC-BY 4.0)

According to the data presented by Hozak et al. (1993),^92^ the bodies containing newly synthesized DNA (replisomes) are fixed to the fibril elements of nuclear matrix network. Similar data for splicisomes were presented by Wan et al. (1999).^95^

In support of the dana on the role of RNA in stability of the internal nuclear matrix obtained on the ultrathin sections (Herman et al., 1978; Smith et al., 1984),^9,13^ it was shown using thick preparations of residual nuclei that RNase makes soluble the fibrogranular network forming internal nuclear matrix while the peripheral fibrils forming the lamina are saved (Fey et al., 1986; Hozak et al., 1995).

Consequently, the internal nuclear matrix IS distinguished by dependence of its integrity on RNA. Usually, nuclear RNA is present in complex with certain proteins, that is, in the form of a ribonucleoprotein (RNP). He et al. (1991)^96^ using immuno electron microscopy have found that the protein belonging to RNP, and therefore RNP itself, are present in the granular components of the internal nuclear matrix, but not in the fibrils. It means that destroying of granules ruins the whole network making it soluble. However, according to other data obtained using similar methods, RNP is present in the fibrils of the internal nuclear matrix (see Nickerson, 2022)^93^. On the other hand, several studies using immunological methods have shown the presence of lamin A in the internal nuclear matrix (Hozak et al., 1995; Neri et al. 1999a,b).^81,82,97^ Wether RNP and lamin A form separate fibrils, or they are present in the same fibril, has not yet been determined. It should be noted that many separate lamin A molecules are, probably, present in nucleoplasm (Naetar et al., 2017; Danielsson et al., 2023).^98,99^

Here an interesting problem arises. Lamin A/C gene is activated only when cell begins to differentiate (<u>Constantinescu</u> et al., 2006).^100^ Therefore, embryonic stem cells are devoid of lamins A and C. Composition of nuclear matrix fibrils in these cells is of interest.

Almost always, experiments on two-dimensional electrophoresis of nuclear matrix proteins show the presence of nuclear actin. However, its functions have been unknown for a long time. Later, many functions of nuclear actin were proposed. Thus, according to Baarlink et al. (2017),^101^ a temporary network of nuclear F-actin is formed in the nuclei newly formed in the daughter cells after mitosis. This F-actin network facilitates the transition from condensed chromosomes to interphase chromatin.

A lot of data has been obtained indicating the participation of actin in the organization of interphase chromatin structure and regulation of gene activity (Shumaker et al., 2003; Visa and Percipalle, 2010; Moore and Vartiainen, 2017; Xie and Percipalle, 2018; Miyamoto and Harata, 2021; Mahmood et al.,2022).^102,103,104.105,106,107^ More details are necessary for describing the mechanism of nuclear actin activities.

Nuclear bodies are still one component of nucleoplasm. The nucleolus is classified as a nuclear body. However, most nuclear bodies are small particles of 0.1-1 µm in size. The basis of such a body is a long non-coding RNA molecule (lncRNA) (Kawaguchi et al., 2015; Nakagawa et al., 2022).^108,109^ Depending on the structure of the lncRNA, a set of corresponding proteins is attached. The membrane does not form (Mao et al., 2011; Yamazaki et al., 2019),^110,111^ and therefore the nuclear bodies are ready for interactions.

Nuclear bodies containing the protein PML called “PML nuclear bodies” are bound to the nuclear matrix (Lallemand-Breitenbach and Thé, 2010; Silonov et al., 2023).^112,113^ They participate in DNA replication, reparation, transcription, RNA splicing (Chang et al., 2018; Silonov et al., 2023).^113,114^ Perhaps the PML corpuscles serve as platforms on which enzyme complexes that carry out and regulate genetic processes are assembled.

An important problem is the fate of the components of the nuclear matrix during mitosis. Laemmli and his colleagues have shown that DNA loops are retained in condensed mitotic chromosomes (Mirkovitch et al., 1988). ^35^ Consequently, the elements of the nuclear matrix that form these loops are also retained in mitosis. Moreover, they should double when the sister chromatids begin to diverge. Proteome analysis showed that 67% of proteins present in interphase nuclear matrix are found also in mitotic chromosome scaffold (Sureka et al., 2018).^115^

It is believed that this particular scaffold plus also retained nuclear bodies are the seeds that preserve the “mitotic memory” and reproduce the architecture of the maternal interphase nucleus in the newly formed daughter nuclei. In particular, decondensed chromosomes occupy their territories, and their conformation, which determines which genes are active, becomes characteristic of a certain type of cell (Nickerson and Penman, 1992; Sureka et al., 2018; Soujanya et al., 2023).^115,116,117^

All the above descriptions and discussions were based on a nuclear matrix model consisting of an peripheral lamina and an internal nuclear matrix. Moreover, according to the experiments involving autoradiography and immunochemistry, most of the DNA fixation sites are localized in the internal nuclear matrix, as well as the enzyme complexes that carry out DNA replication, transcription and splicing (Herman et al., 1978; Smith et al., 1984; Hozak et al., 1993, 1994,.1995; Hozak, 1996; Wei et al., 1999; Bihani et al., 2023).^8,9,13,82,92, 118,119,120^

However, it should be noted that since the 70-s of the last century, another, fundamentally different idea of the structure of the nuclear matrix has been proposed. After chromatin removal, Blobel and co-authors observed residual nuclei in the form of a ring formed by nuclear pores-lamina with an empty inner space (Aaronson and Blobel, 1975; DwvER. and G. BLOBBL 1976; Gerace and Blobel, 1980; Gerace and Huber, 2012; Almendáriz-Palacios et al., 2020).^63,121,122,123,124^ According to this concept, the bases of DNA loops are fixed in the lamina, that is, on the periphery of the nucleus, and the loops themselves are directed into the “empty” interior of the nucleus, which turns out to be filled with chromatin.

Many authors have adopted this idea of the cell nucleus architecture and used it to interpret their experimental results (Hutchison et al., 1994; Tajik et al., 2016; Turgay et all., 2017; Sapra and Medalia, 2021; Kim et al., 2023; Madsen-Østerbye et al., 2023; Pennarun et al., 2023).^54,79,80,125,126,127,128^

Proponents of this model of cell nucleus architecture believe that the internal nuclear matrix is an artificial structure formed by aggregates of proteins and RNA during the production of residual nuclei under non-physiological conditions, especially when using high ionic (Singer et al., 1997).^129^ According to Tan et al. (2000)^130^ proteins of the RNP complexes released due to exposure to RNase or high ionic strength form long fibrils and networks. Visually these artificial constructions are similar to those considered as an internal nuclear matrix. However, RNase actually removes the internal nuclear matrix, rather than creating it (Herman et al., 1978; Smith et al., 1984; Fey et al., 1986; He et al., 1990; Nickerson, 2022).^9,13,90,91,93^ Several authors used chromatin removal methods that excluded extraction with a high salt medium. Nevertheless, they observed a fibrillar-granular network of the internal nuclear matrix (Jackson and Cook, 1988; Hozak et al.,1993; Wan et al., 1999).^92,94,95^ The above-described experimental results with [^3^H] or Fluorescent labeled nucleic acids and immunochemically labeled proteins participating in the genetic processes have directly shown that sites of DNA fixation, nascent (puls labeled) RNA and DNA, enzyme complexes that carry out DNA replication, transcription and splicing are located mainly in the internal regions of the nucleus (see above). These results, obtained mainly at the end of the last century, have not yet been disputed.

Many details of the composition and functions of the internal nuclear matrix are already known and have been described above.

However some important details are not yet elucidated.

Thus, the presence of the fibrillar nuclear protein lamin A in the filaments of internal nuclear matrix (Hozak et al., 1995)^82^ was expected, but the crucial role of RNA in maintaining the integrity of these filaments or their release from an insoluble network shown in several studies (Herman et al., 1978; Smith et al., 1984; Fey et al., 1986)^9,13,90^ was not expected. The exact location of the nuclear matrix RNA (probably, a long non-coding RNA) has not yet been discovered.

In addition to lamin A, actin and proteins that fix DNA in matrix, many other proteins were detected in the internal nuclear matrix using a two-dimensional PAGE (Berezney and Coffey, 1977;. Gerace and Blobel, 1980; <u>Peters</u> and <u>Commings</u>, 1980; <u>Kaufmann</u>, 1989; Cupo et al., 1990; Ma et al., 1999; Bihani et al., 2023).^58,60,61,62,63,120,131^ Most of them have not yet been identified. Consequently, some structures and functions of the nuclear matrix have to be explored.

In order to perform its very precise functions, which allow genetic processes to take place, the nuclear matrix must have a highly organized structure. However, so far, no ordered nuclear matrix has been observed in mammals. The only communication on highly ordered structure of nuclear matrix in *Xenopus* oocytes was from Aebi et al. (1986).^76^ But these data have not been confirmed by similar data from other authors.

Still one problem is evident.

The network of the internal nuclear matrix, observed by various authors in an electron microscope, consists of very thin (about 10 nm) and, consequently, mechanically weak filaments (Capco et al., 1983; Fey et al., 1986 ; Jackson and Cook, 1988; He et al., 1990; Hozak et al., 1993; Nickerson, 2022).^88,90,91,92,93,94^

However, the nuclear matrix performs some important mechanical functions: organizing the architecture of the nuclear interior, controlling the correct location of chromatin, holding the nucleoli in their places. Besides, the elements of the nuclear matrix that fix the bases of chromatin loops are preserved in condensed chromosomes during mitosis. They control chromatin conformation in these chromosomes and are responsible for their elongated form.

For the fulfillment of all these functions, nuclear matrix must have some rigid and durable components, which the described thin filaments cannot be. It can be assumed that some of its components were lost during the preparation of the nuclear matrix. Therefore, the study of the nuclear matrix using various approaches should be continued.

Here cross-striated (axial repeat about 65 nm) thick (30-80 nm) linear (non-branching) nuclear fibrils are described. Externally, they resemble the collagen fibrils of extracellular matrix. Therefore, the main task of the experimental part of this article was to prove the localization of the detected fibrils in the cell nucleus.

## EXPERIMENTAL PROCEDURES

### Isolation of cell nuclei

The nuclei were isolated from calf and rat tissues.

The calf’s tissues were purchased at an abattoir and immediately frozen in liquid nitrogen. They were thawed immediately before the experiment. Rat tissues (Wister males) were isolated before the experiment.

The tissue was cut into small pieces and homogenized by 5-7 strokes in loosely fitted Dounce hand-homogenizer in the presence of 3 volumes of “complete medium” (CM)**: 2**0 mM Tris-HCl (pH 7.5), 100 mM KCl, 50 mM NaCl, 1 mM MgCl_2_, 1 mM CaCl_2_, 0.2 mM PMSF. The homogenate was filtered through six layers of cheesecloth and centrifuged at 700 g for 10 min. The pellet was suspended in CM, and Triton X-100 was added to a final concentration of 0.5%. After exposure at 0°C for 10 min, the suspension was layered on the top of 5% sucrose in CM and centrifuged at 1000 g for 15 min. Pelleted nuclei were carefully suspended in CM, and the centrifugation was repeated. Peleted nuclei were suspended in CM at 30-50 A_260_/ml layered on the top of 2.2 M sucrose in CM and centrifuged in the swinging-buckеt rotor at 100000 g for 120 min. White pellet of purified nuclei was suspended in CM at 30-50 A_260_/ml. All the procedures were carried out at 0-+4°C.

### Mechanical destruction of the cell nuclei

The cell nuclei suspended in CM were treated with ultrasound in the tube striktor by 15 sec pulses at 30 wt (I=0.5A) and 22 kHz. During,exposure to ultrasound the temperature of the suspension was 6-8°C.

### Isolation of nucleoli

The preparations containing isolated nucleoli were obtained mainly according to Peters and Comings (1980)^58^ and Hacot et al. (2010). Suspended in CM cell nuclei isolated from rat liver (10^5^ nuclei/ml) were treated with DNase I (10 µg/ml) at 20°C for 15 min and treated with ultrasound as described above for 120 sec. The sonicate was layered on the top of CM containing 5% sucrose and centrifuged in swinging backed rotor at 3000 g for 10 min. The pellet of the nucleoli was suspended in CM and the centrifugation was repeated. The pellet of the purified nucleoli was suspended in CM and studied in electron microscope as whole mounts.

### Preparation of residual nuclei

Two variants of partially extracted (residual) nuclei were used. According to the first of them the nuclei suspended in CM were treated with 1% sodium deoxycholate and 1% non-ionic detergent Tween 40 at 0°C for 40 sec, immediately layered on the top of 5% sucrose in CM supplied with 2 M NaCl and centrifuged at 1500 g for 15 min at 0-4°C. The residual nuclei (1-st variant) were suspended in CM at 30-50 A_260_/ml and studied in electron microscope.

For more thorough removal of chromatin, these residual nuclei were subjected to further treatments: DNase I and RNAase A (20 µg/ml each) were added, and the mixture was incubated at 20°C for 10 min and at 0°C for 20 min. After this, Triton X-100 was added to 0.5%, and the incubation was continued at 0°C for 5 min. Then the suspension was layered still ones on the top of 5% sucrose in CM and centrifuged at 3000 g for 20 min. The pellet of residual nuclei (2-nd variant) was suspended in CM and studied in electron microscope.

### Electron microscopy

## RESULTS

### Observation of isolated from calf and rat tissues and highly purified cell nuclei. Finding of cross-striated fibrils associated with damaged nuclei

Figure 1 shows typical examples of cell nuclei isolated from calf and rat tissues. They are well purified from other tissue and cell components (1A,1B and 1C). However, in these preparations of isolated nuclei, rare nuclei can be observed that look damaged. Cross-striated fibrils are located in close contact with these nuclei. It looks as if the fibrils are released from the nucleus through a damage in its envelope (1D,1E,1F and 1G).

**Figure 1.**
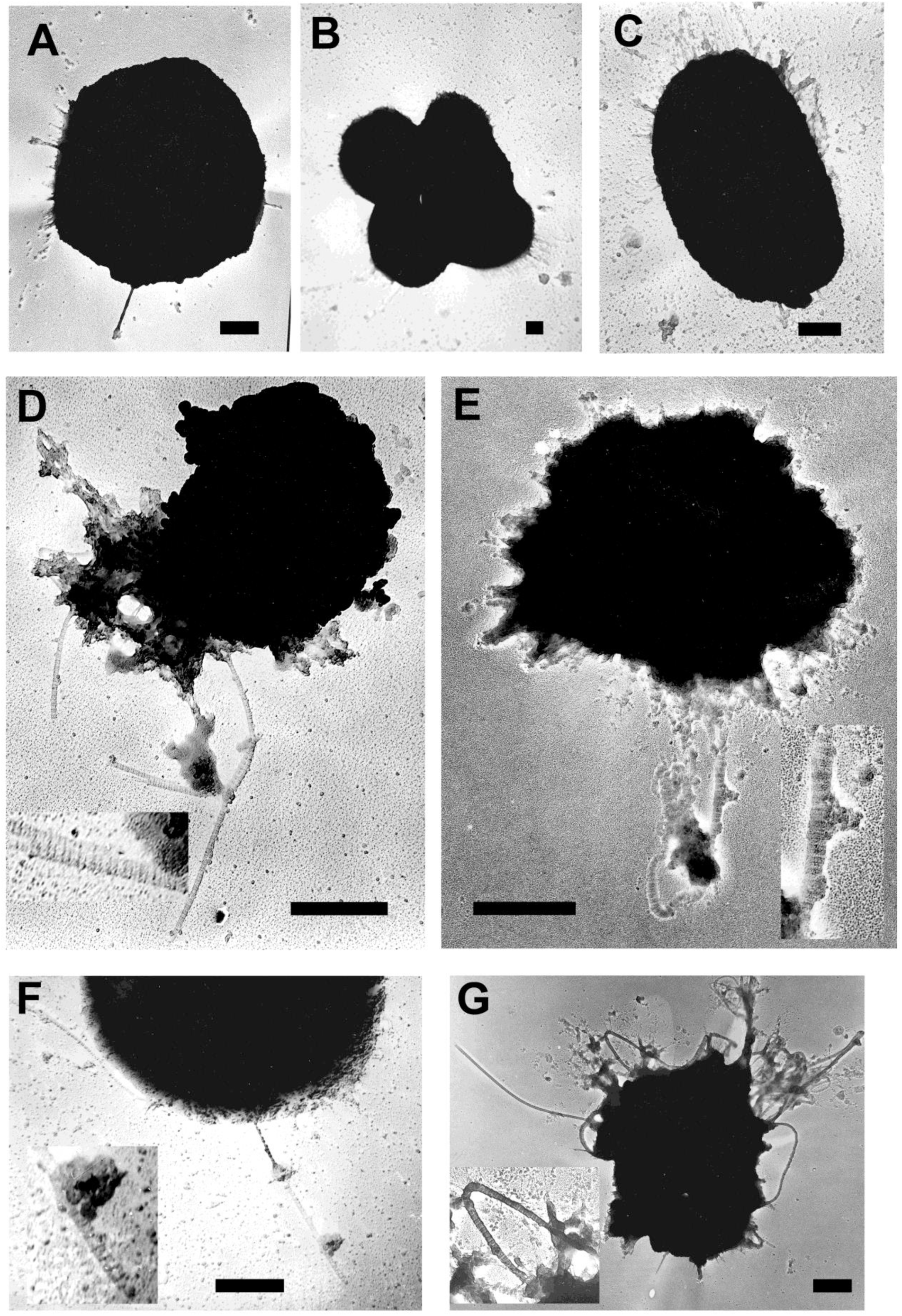
Highly purified cell nuclei. The nuclei isolated from calf liver (A), from rat liver (B), from rat spleen (C). Damaged nuclei observed in the same preparations: from calf liver (D,E), from rat liver (F,G). Shadowing with Pt-Pd alloy. Bar 1 micron

### Observation of thick cross-striated fibrils in the material of destroyed by ultrasound cell nuclei isolated from the calf and rat tissues

Ultrasound treatment of calf and rat tissue cell nuclei with short pulses for 60 seconds (see Experimental procedures) destroys them and reveals the same cross-striated fibrils that are clearly visible on the free film, as well as a granular substance, presumably chromatin (Figure 2A 2B and 2 C).

**Figure 2.**
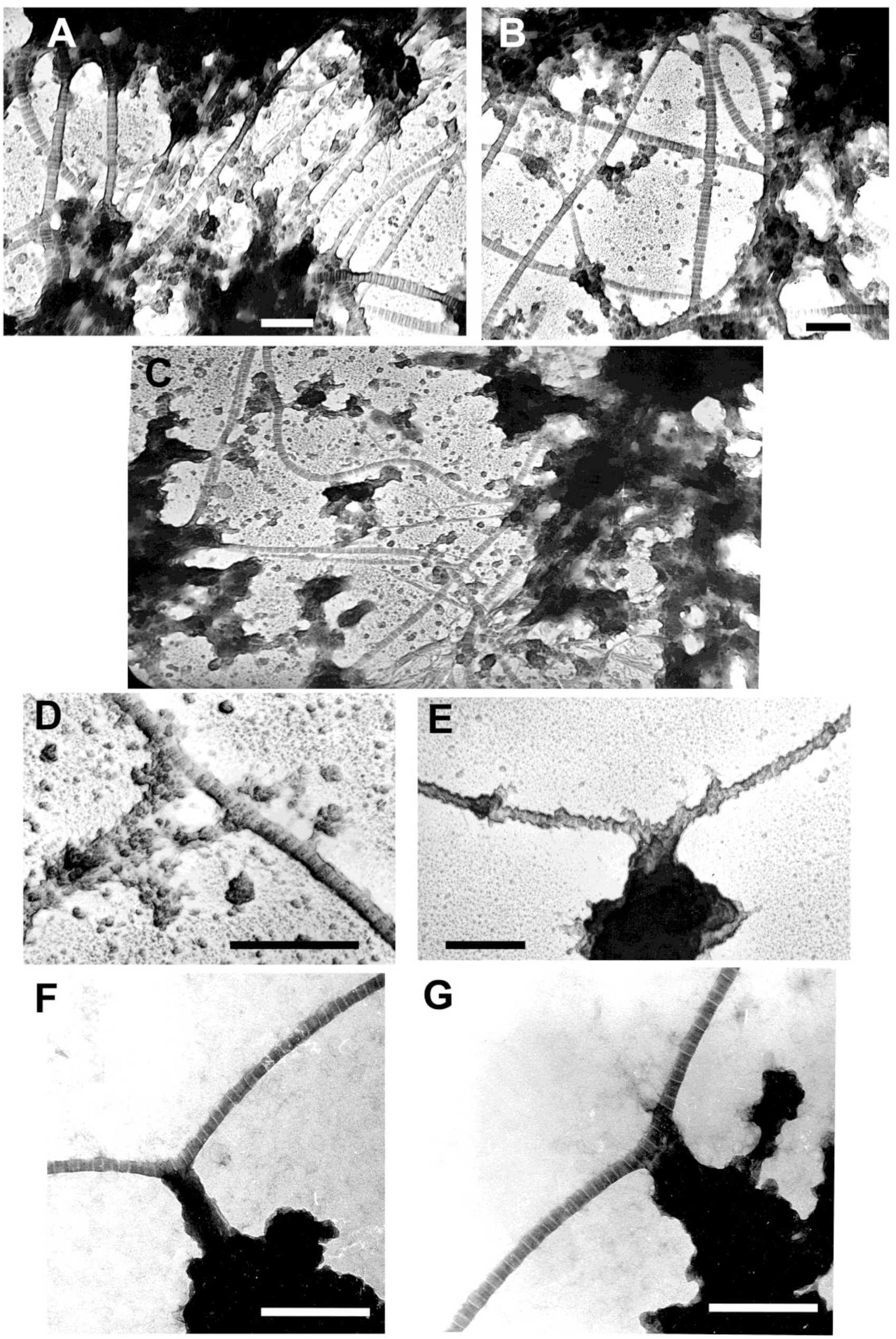
Material of cell nuclei disintegrated by ultrasound. (A) Calf liver. (B) Rat liver. (C) Rat spleen. (D,E,F,G) Regions of tight contact between thick cross-striated fibrils and granular substance. (D,E) Calf liver. (F,G) Rat liver. (A,B,C,D) shadowing with Pt-Pd alloy. (E,F,G) staining with uranyl acetate. Bars 0.5 micron

In preparations of destroyed cell nuclei shown in Figure 2, the sites of tight contact (connections) of cross-striated fibrils with granular material are present (Figure 2D,2E,2F and 2G).

When the nuclei before ultrasound treatment were treated with micrococcal nuclease (30 µg/ml, at 37°C for 15 min), there was no condensed granular substance in the preparations. Instead, closely spaced small particles (residual nucleosomes?) and the same cross-striated fibrils were observed (Figure 3). This result confirms the assumption that the granular material observed in preparations of destroyed nuclei shown in Fig. 3 is indeed chromatin.

**Figure 3.**
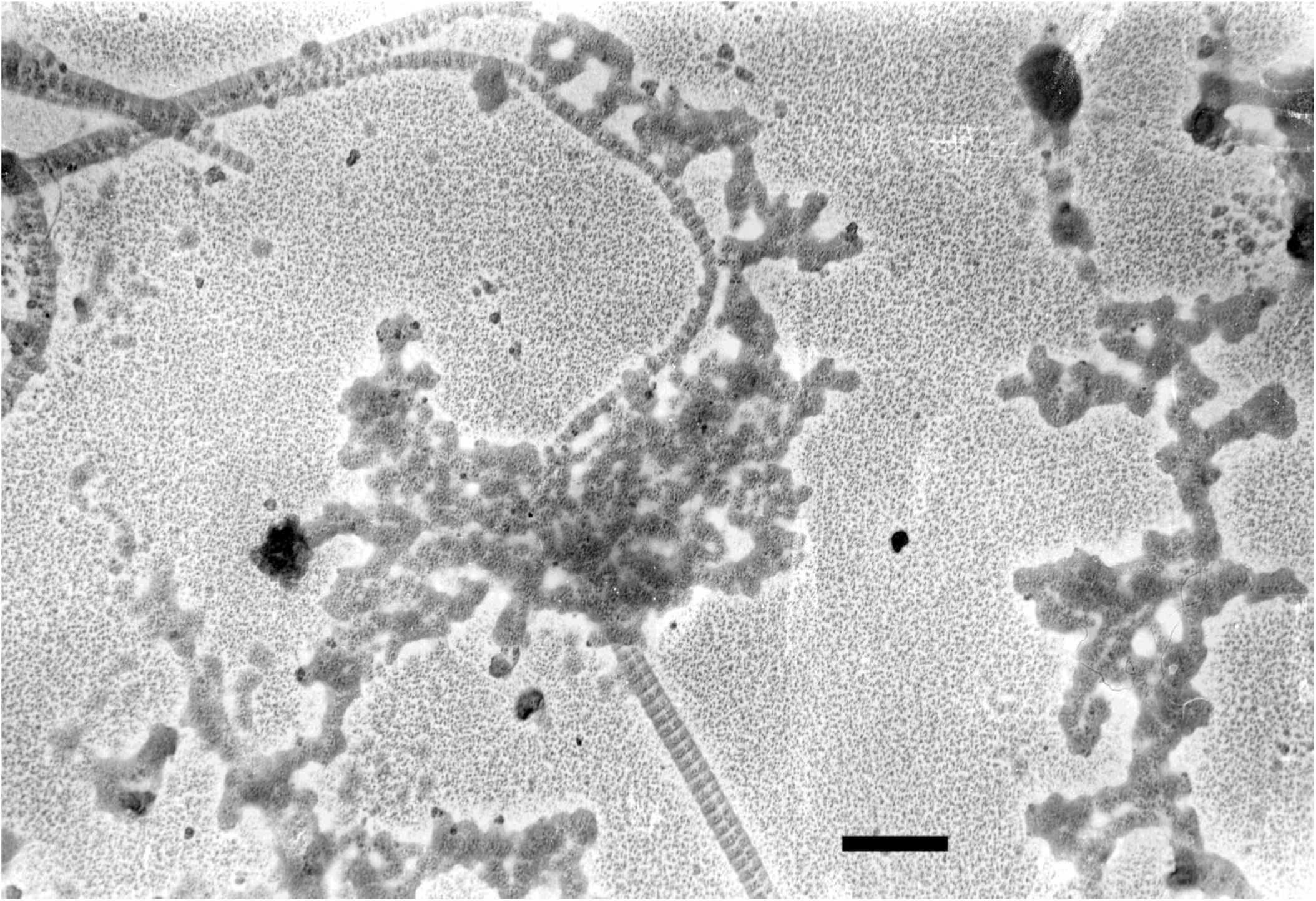
Material of disintegrated cell nuclei of rat spleen limitedly treated with Micrococcal nuclease (30 microgram/ml, 37°C, 10 minutes). Shadowing with Pt-Pd alloy. Bar 0.5 micron

### Observation of cross-striated fibrils associated with isolated nucleoli

It has been observed that the cross-striated fibrils released from damaged nuclei are sometimes associated with nucleoli-like bodies (see Figure 2). Therefore, nucleoli were isolated from the purified rat liver nuclei. (see Experimental procedures). Many of these nuclear particles proved to be connected with cross-striated fibrils (Fig. 4). In some cases, it can be seen that these fibrils pass through the nucleoli. It is likely that heavy nucleoli are held in place in the nucleus precisely because they are connected to thick probably quite durable fibrils.

**Figure 4.**
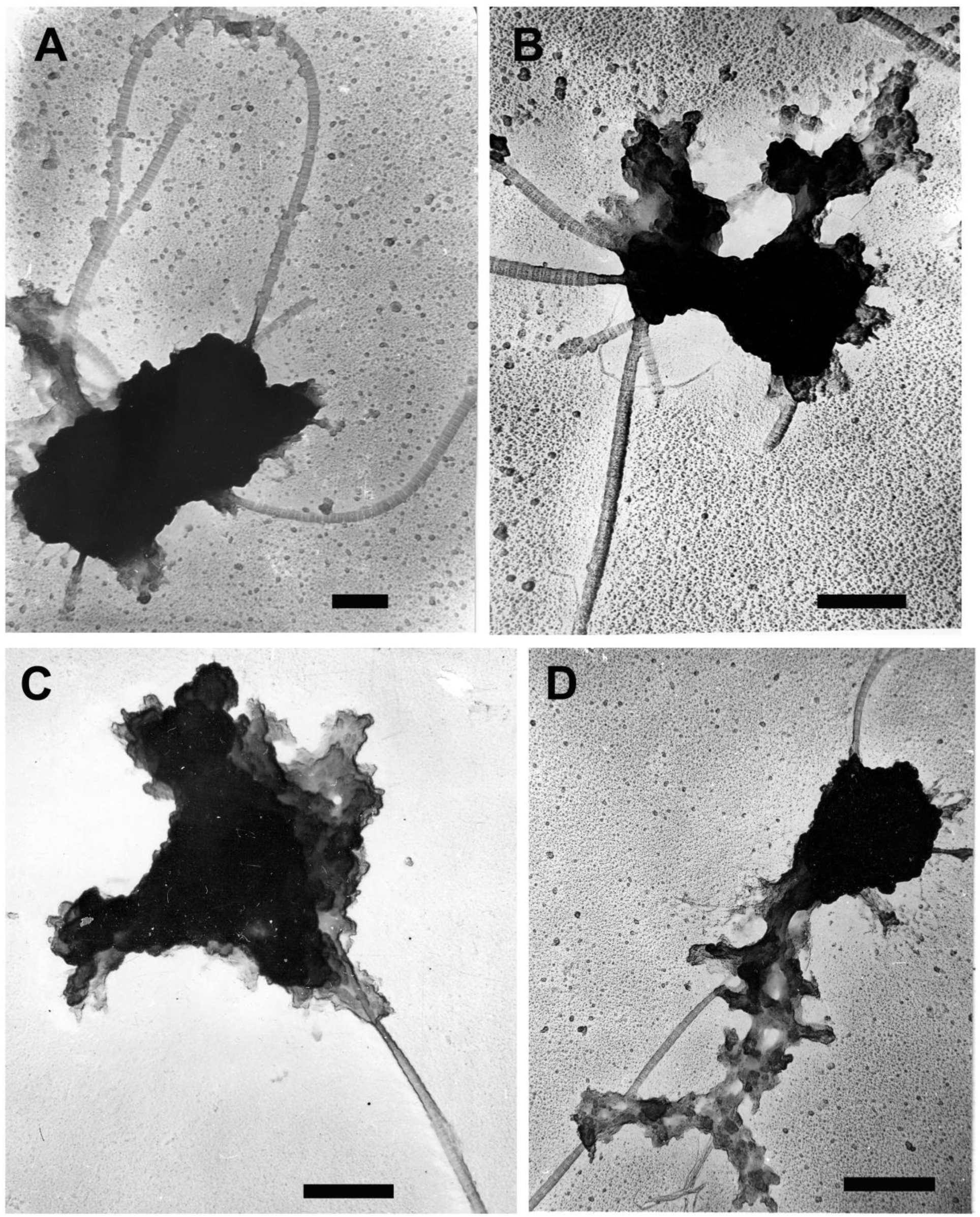
Nucleoli connected with thick cross-striated fibrils obtained from the cell nuclei isolated from calf liver. Shadowing with Pt-Pd alloy. Bars 0.5 micron

### Observation of fibrillar structures in residual nuclei

Treatment of isolated cell nuclei with deoxycholate and Tween 40 (1-st variant of residual nuclei, see Experimental procedures) leads to the formation of ruptures in the nuclear envelope through which the cross-striated fibrils are released (Figure 5A,B). Through these cracks a great amount of the nuclear material is thrown out, and some nuclei become partially transparent in electron microscope. Due to this, it is possible to observe some details of the nuclear structure in the “whole mount” preparations. In particular, nuclear pores become visible (Fig. 5A). It is possible also to recognize cross-striated fibrils passing near the nuclear periphery (Fig. 5A). The presence of nuclear pores definitely confirms that the entire object is a cell nucleus, and the observed fibrils are indeed nuclear components.

**Figure 5.**
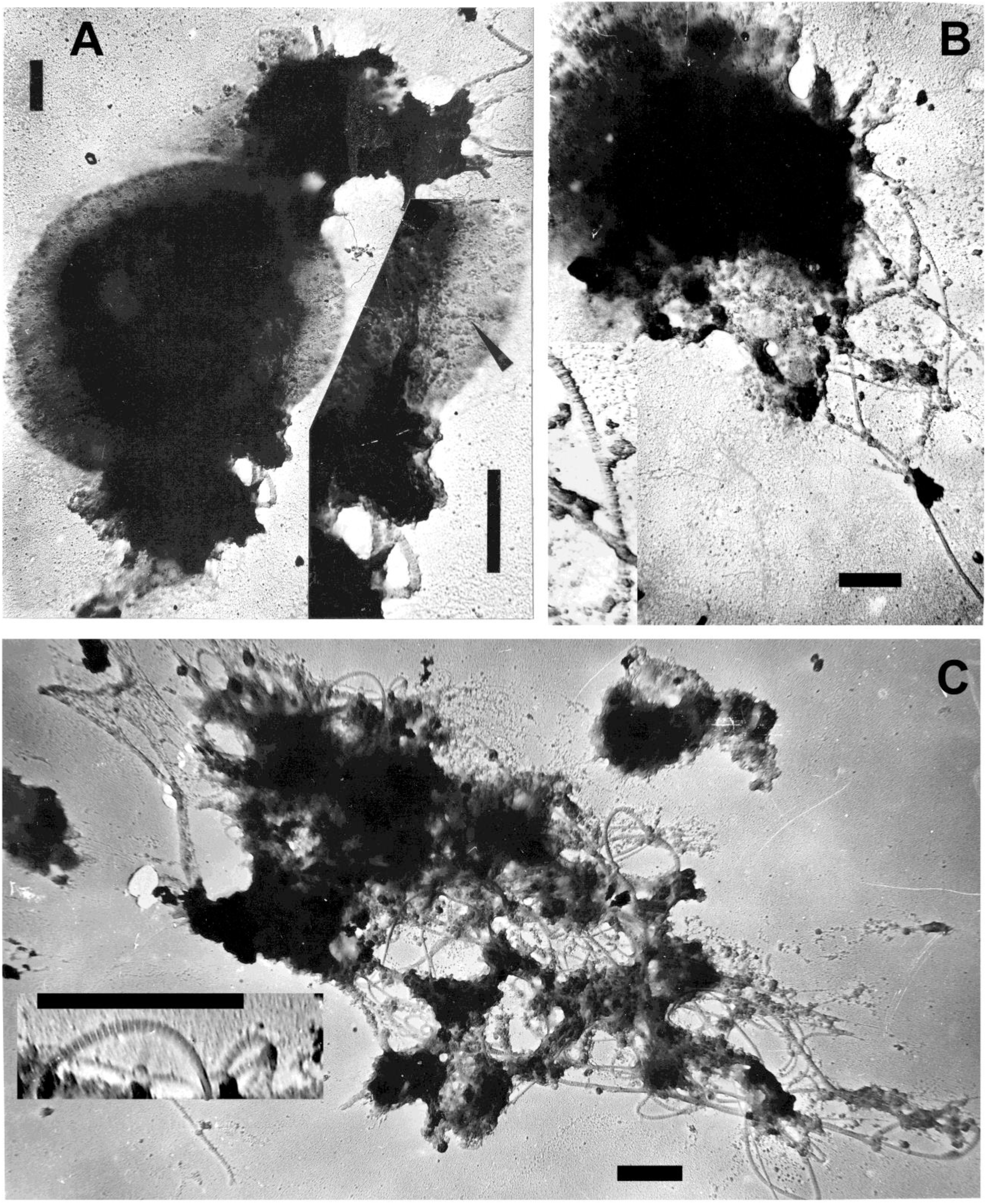
Residual nuclclei. (A,B) Residual nuclei obtained by the treatment of the cell nuclei from rat liver with 1% desoxicholate 1% Tween 40 (1-st variant). (C) These residual nuclei additionally treated with 2 M NaCl, nucleases and 0.5% Triton X-100 (2-nd variant). Figure 5. Residual nuclclei. (A,B) Residual nuclei obtained by the treatment of the cell nuclei from rat liver with 1% desoxicholate 1% Tween 40 (1-st variant.). (C) These residual nuclei additionally treated with 2 M NaCl,nucleases and 0.5% Triton X-100 (2-nd variant). Shadowing with Pt-Pd alloy. Bars 1 micron

Additional treatment of the nuclei with 2 M NaCl, nucleases and Triton X-100 (2-nd variant of residual nuclei, see Experimental procedures) leads to loss by them of chromatin and of significant part of the nuclear envelope. As a result, it became possible to clearly observe the internuclear network of thin filaments (Figure 5C). Thick cross-striated fibrils are also seen in this preparation.

In other samples of 2-nd variant, elements of the nuclear matrix are visible, where thin filaments are connected to thick cross-striated fibrils (Figure 6).

**Figure 6.**
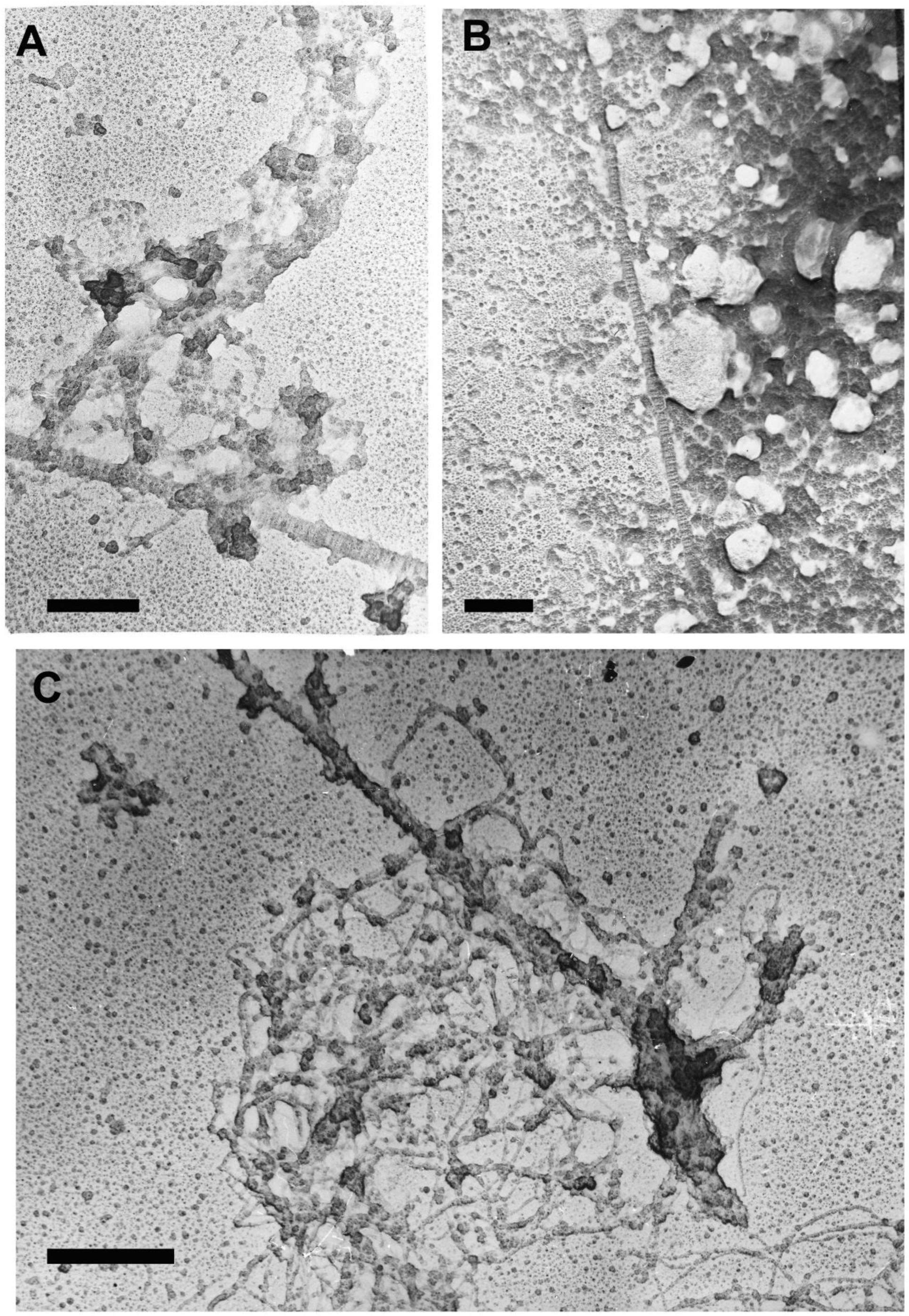
Associations of the nuclear matrix network with thick cross-striated fibrils observed in the preparations of the residual nuclei (2-nd variant) from rat liver. Shadowing with Pt-Pd alloy. Bars 0.5 micron

### Cross-striated fibrils as axial elements in condensed chromosomes in mitosis

In the organs containing proliferating cells (thymus, spleen, regenerating liver etc.), some of these cells are in prophase of mitosis. Their nuclei contain condensed chromosomes. The material of cell nuclei, which were isolated from these organs and treated with ultrasound for 30 seconds, contained dense elongated bodies with bundles of thick cross-striated fibrils as the axial elements (Figure 7A,B). Presumably, these structures could be fragments of prophase chromosomes damaged by ultrasound. Indeed, pretreatment of the nuclei with DNase I (10 µg /ml, 20°C, 10 min) led to the appearance of less dense domains in these bodies (Figure 7C), which indicates the presence of DNA and, therefore, confirms that these bodies are fragments of prophase chromosomes.

**Figure 7.**
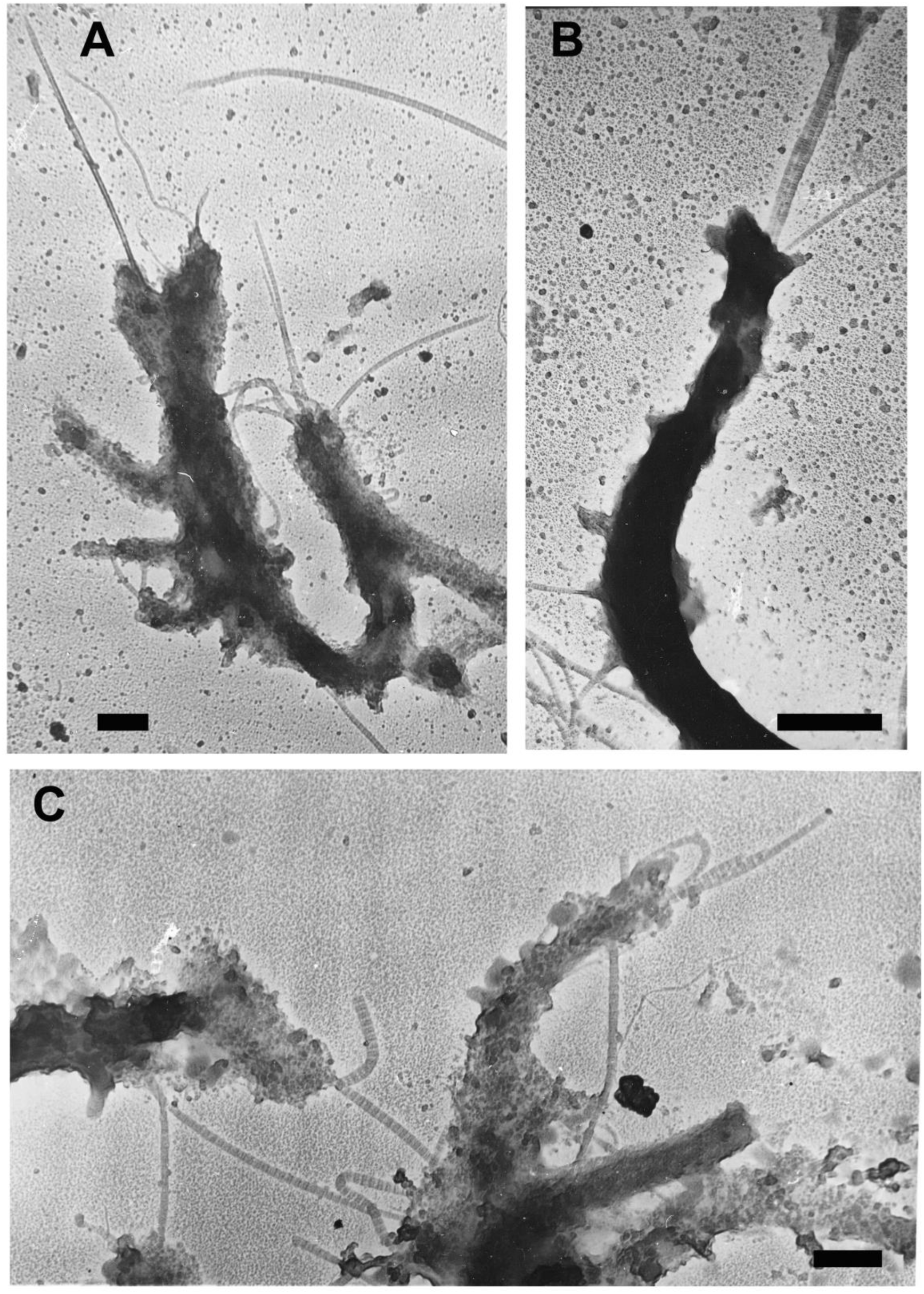
Preservation of thick cross-striated fibrils in mitosis. (A,B) Dense elongated bodies with the bundles of thick cross-striated fibrils as axial elements observed in the material of treated with ultrasound rat spleen nuclei, some of which contain prophase chromosomes. (C) Less dense (dispersed) elongated bodies, when the nuclei preliminarily were treated with micrococcal nuclease, what conforms that these bodies are fragments of prophase chromosomes. Shadowing with Pt-Pd alloy. Bars 0.5 micron

### Some properties of thick cross-striated nuclear fibrils

Thick cross-striated nuclear fibrils (axial repeat D is about 65 nm) do not have branches and have the same thickness along their entire length, but the thickness of different fibrils varies from about 40 to 100 nm. Thick cross-striated nuclear fibrils are stable in the “complete medium” (CM). The salt composition of this medium mimics the physiological salt composition of the cell nucleus (<u>Paine</u> et al., 1981).^132^ These fibrils remain stable at elevated ionic strength (in 2 M NaCl), but at low ionic strength (10 mM Tris-HCl buffer, pH 7.4) they are unstable. Proteolytic enzymes chymotrypsin and collagenase destroy thick cross-striated fibrils (data not shown).

## Discussion

The data presented in the Results show that thick cross-striated linear (non-branching) fibrils are present in the cell nuclei of mammalian tissues. They probably form a firm skeleton of the nuclear matrix, which also contains a network of thin filaments that can be seen in the residual nuclei (Figure. 5c).

Interestingly, thin filaments were practically not observed in the preparations treated with ultrasound (see Figure 3). Perhaps, after the destruction of an insoluble network of thin filaments due to this treatment, they become free and can be removed during the washing of preparations on water drops before electron microscopy (see Experimental procedures).

In preparations of nuclei destroyed by ultrasound, many sites of close contact between chromatin and thick cross-striated fibrils were found (Figure 3C-F). Some of these contacts can be accidental, but real connections also exist. These contacts can be so strong that they locally distort the structure of the fibrils (see Figure 3D-F). The presence of these natural contacts shows that thick cross-striated fibrils are involved in the control of chromatin conformation and placement inside the nucleus. In particular, cross-striated fibrils are associated with the nucleoli (Figure 5). Probably, these fibrils fix the position of massive nucleoli in the space of the nucleus.

It is probable also, that cross-striated fibrils participate in till now mysterious process of condensed chromosomes formation in mitosis. Rearrangement of these fibrils can lead to redistribution of interphase chromatin into condensed chromosomes. The fibrils themselves occupy the position of axial elements in these chromosomes, which provide the linear shape of the chromosome (see Fig. 6). As axial elements of chromosomes, thick cross-striated fibrils are preserved during cell division. With them their contacts with chromatin are preserved. Between these contacts, contacts with special DNA regions that form the loop conformation of chromatin can be preserved. This would mean that thick cross-striated fibrils ensure the preservation of chromatin conformation in the form of loops over generations.

In this work, various experimental approaches were used to prove the presence of thick cross-striated fibrils in the cell nucleus, the results of which were analyzed using an electron microscope (see the Results)

The problem is complicated by the fact that the axial repeat (D) in these fibrils (about 65 nm) is similar to that in collagen fibrils of the extracellular matrix, which are formed in the extracellular medium using a secreted collagen precursor (Kadler, 2017; Holmes et al., 2018).^133,134^ So far, such fibrils have not been found either in the cell nucleus or in the cytoplasm.

An obvious question arises why many authors who used similar approaches to the isolation of the cell nuclei and to the study of the nuclear matrix did not observe the thick linear cross-striated fibrils described here. According to their observations, the nuclear matrix is organized as a network of thin (about 10 nm) branched filaments. The probable reason for the absence of thick linear cross-striated fibrils, at least in some cases, can be explained by a too long Staying of the nuclear matrix preparations in a buffer of low ionic strength, since these fibrils are destroyed in such an environment. Perhaps there are other reasons that can be voiced in the course of subsequent discussions.

The results presented in this paper ascribe to thick linear cross-striated nuclear fibrils crucial role in nuclear architecture, chromatin conformation and other phenomena that should be studied in special experiments.

However, first of all, it is necessary to confirm these results by the data of other authors. This will serve as an incentive for further research, in particular, the isolation and purification of thick cross-striated nuclear fibrils and the clarification of their composition by the use of mass spectrometry etc.

## Supporting information

Supplemental Images

## Acknowledgments

I am grateful to Vera A. Novitskaya and Galina Yu. Skladtchikova for help in preparation of purified cell nuclei and to Olga M. Gorbenko for preparing illustrations for publication. This work was supported by the Institute of Macromolecular Compounds and the Petersburg Nuclear Physics Institute named by В.Р.Konstantinof

