## Supplementary figures and images for "Detection and electron microscopic study of thick cross-striated linear fibrils in mammalian cell nuclei"

### Supplemental Images

**Supplement**

Large-scale images

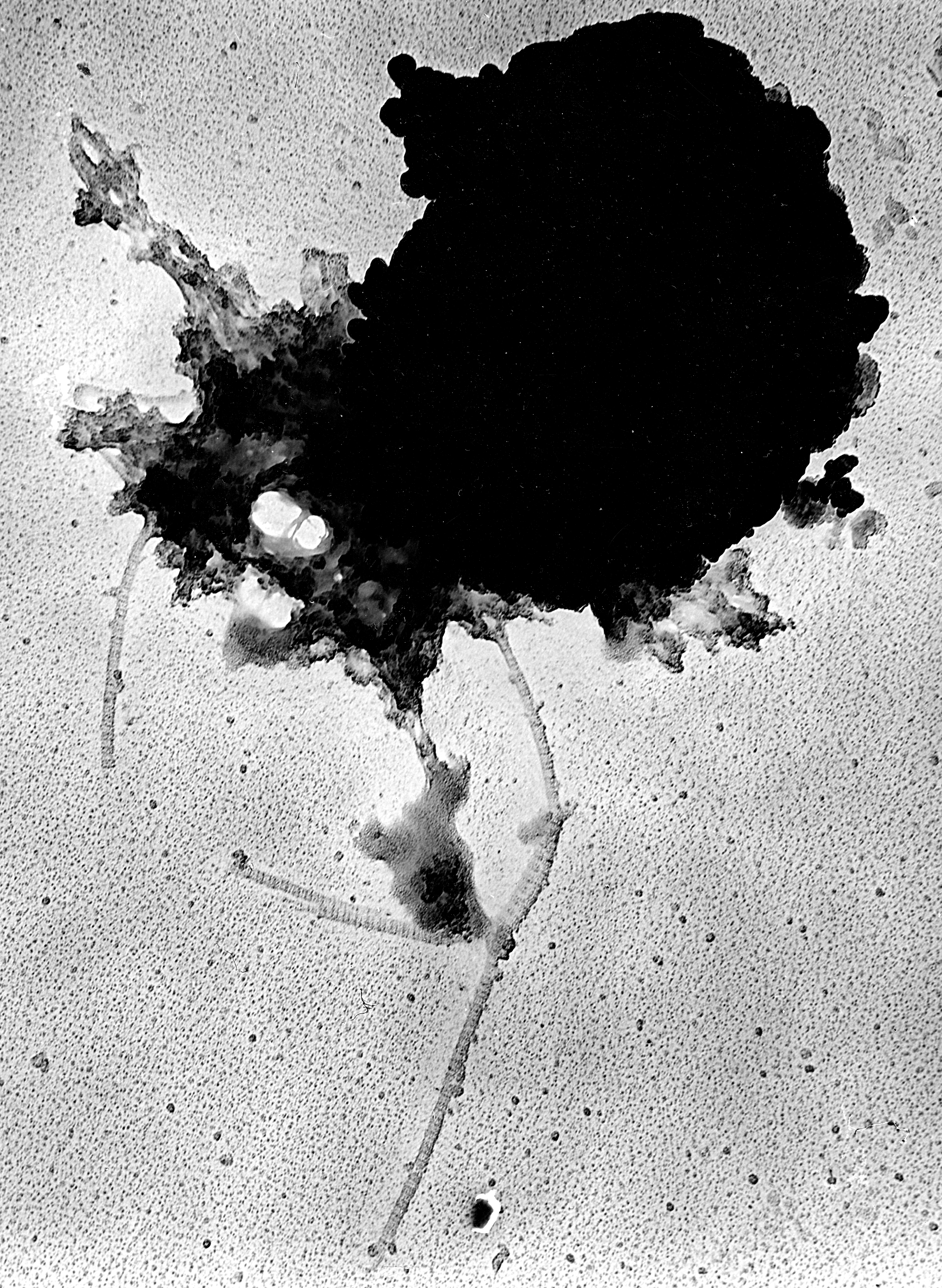

Figure 1D(S). Bar 1 micron

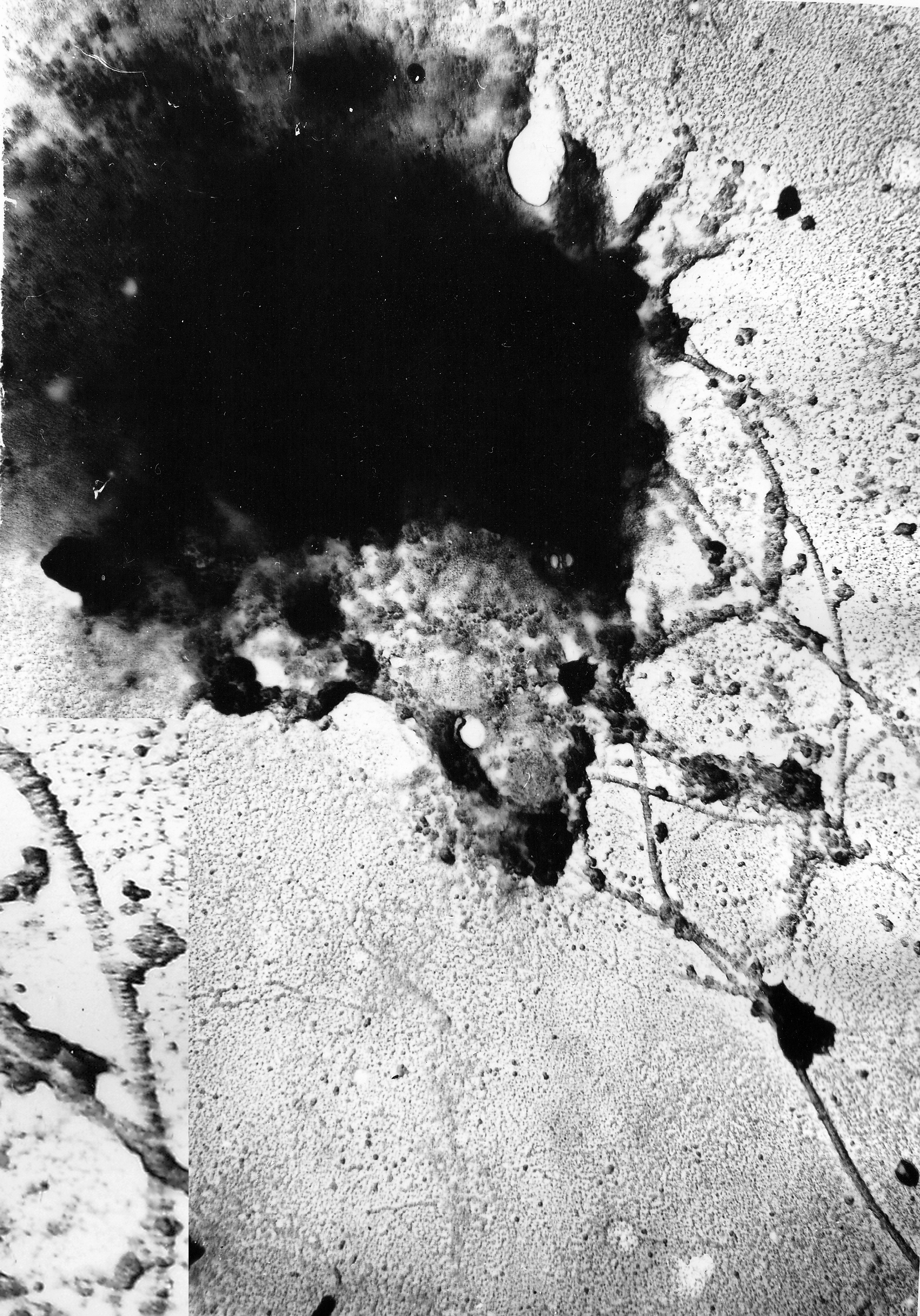

Figure 5B(S). Bar 1 micron
